# Art2 mediates selective endocytosis of methionine transporters during adaptation to sphingolipid depletion

**DOI:** 10.1101/2022.05.20.492841

**Authors:** Nathaniel L. Hepowit, Bradley Moon, Robert C. Dickson, Jason A. MacGurn

## Abstract

Accumulating evidence in several model organisms indicates that reduced sphingolipid biosynthesis promotes longevity, although underlying mechanisms remain unclear. In yeast, sphingolipid depletion induces a state resembling amino acid restriction, which we hypothesized may be due to altered stability of amino acid transporters at the plasma membrane. To test this, we measured surface abundance for a diverse panel of membrane proteins in the presence of myriocin, a sphingolipid biosynthesis inhibitor. Unexpectedly, we found that surface levels of most proteins examined were either unaffected or increased during myriocin treatment, consistent with an observed decrease in bulk endocytosis. In contrast, sphingolipid depletion triggered selective endocytosis of the methionine transporter Mup1. Unlike methionine-induced Mup1 endocytosis, myriocin triggers Mup1 endocytosis that requires the Rsp5 adaptor Art2, C-terminal lysine residues, and the formation of K63-linked ubiquitin polymers. These findings reveal cellular adaptation to sphingolipid depletion by ubiquitin-mediated remodeling of nutrient transporter composition at the cell surface.

## Introduction

Sphingolipids (SLs) are a diverse class of lipids that serve as a structural component of eukaryotic membranes but also have important regulatory functions related to cell signaling. The first steps of SL biosynthesis occur in the ER and result in the production of ceramide, which is then transported to the Golgi complex for further modification into complex SLs. These complex SLs are then transported to different membranes throughout the cell where they serve a multitude of functions [1]. For example, sphingomyelin regulates sorting of specific secretory cargo in the trans-Golgi network of mammalian cells [2, 3]. In the plasma membrane (PM) of mammalian cells, sphingosine 1-phosphate is generated and can be secreted to act as a signaling molecule that mediates complex processes including vascular development and coordination of immune responses [4]. In yeast, sphingolipids at the PM regulate the activation of TORC2 [5]. Indeed, the variety of regulatory functions served by SLs is underscored by their important role in processes that range from memory and cognition [6] to the progression of cancer [7, 8].

Myriocin (Myr) is a potent inhibitor of serine palmitoyltransferase (SPT) which catalyzes the first step of SL biosynthesis and increases lifespan in a variety of model organisms [9]. There is a growing body of data showing that Myr treatment reduces the severity of age-related diseases in mice and rats, including atherosclerosis and cardiac impairment [10–13], factors for metabolic syndrome, obesity, diabetes and cancer [14–18], amyloid beta and tau hyperphosphorylation in Alzheimer’s disease [19] and other neurodegenerative diseases [20, 21]. Despite its therapeutic potential, it remains unclear how dampening SL biosynthesis confers these health benefits to enhance longevity.

Perturbations that alter sphingolipid homeostasis have complex effects on cellular processes. By characterizing how *Saccharomyces cerevisiae* yeast cells respond and adapt to Myr treatment, we have worked to understand how sphingolipid depletion promotes longevity. Recently, we reported that Myr-treated yeast cells experience a state resembling amino acid restriction, which is associated with decreased uptake of amino acids from the media into the cell [22]. We also reported that Myr triggered the endocytic clearance of the high affinity methionine transporter Mup1 [22]. Based on these prior results, we hypothesized that sphingolipid depletion may trigger broad endocytic clearance of various nutrient transporters. Here, we tested this hypothesis by measuring the surface abundance of a diverse panel of PM proteins – including amino acid transporters, hexose transporters, proton pumps, and signaling receptors. We report that, for most proteins examined, sphingolipid depletion either increases the PM abundance or has no apparent effect on subcellular distribution. Consistent with these observations, we found that Myr inhibits bulk endocytosis while simultaneously triggering selective endocytic clearance of Mup1. We also address the mechanism of Myr-mediated Mup1 endocytosis, which is mechanistically distinct from methionine-induced Mup1 endocytosis. These studies are crucial to understanding how Myr treatment leads to a state of amino acid restriction, and they provide new insights into how the PM is remodeled in response to sphingolipid depletion.

## Results

### Myriocin triggers remodeling of AAT composition at the PM

Our previous finding that Myr treatment generally decreases the intracellular pool of amino acids as well as the rate of amino acid uptake [22] raised the possibility that gradual sphingolipid depletion alters the composition of amino acid transporters (AATs) at the PM. To explore this possibility, we examined cells harboring chromosomal mNeonGreen (mNG) fusions to a panel of AAT genes (*MUP1, CAN1, DIP5, LYP1, BAP2, BAP3, GNP1, HIP1, GAP1*, and *TAT2*) in wildtype (SEY6210) and Vph1-mCherry (a protein localized to the limiting membrane of the vacuole) expressing background strains. Cells at mid-log phase were treated with Myr (400 ng/mL) or mock-treated (solvent) for 5 hours and mixed just prior to visualization by fluorescence microscopy. Vph1-mCherry expression was used to identify cells that had been Myr-treated, while mock treatment was performed on cells lacking any mCherry expression. Quantification of images was performed by measuring mean fluorescence intensity at the PM of individual cells. This analysis revealed that Myr treatment did not significantly alter PM levels of Can1, Dip5, Lyp1, Bap2, Gnp1, or Hip1 (**FIG 1A-B** and **FIG S1A-C**). (In the conditions of this experiment, Gap1 was predominantly vacuole-localized in both mock-treated and Myr-treated cells (**FIG S1A**), and thus localization to the PM could not be quantified.) In contrast, increased levels of Bap3 and Tat2 at the PM were observed in Myr-treated cells (**FIG 1A-B**). Consistent with previously reported results, Myr-treated cells exhibited decreased Mup1-mNG at the PM and increased Mup1-mNG flux into the vacuole (**FIG 1A-B**). Taken together, these results indicate that sphingolipid depletion induces a selective endocytic clearance of the methionine transporter, Mup1.

**Figure 1.**
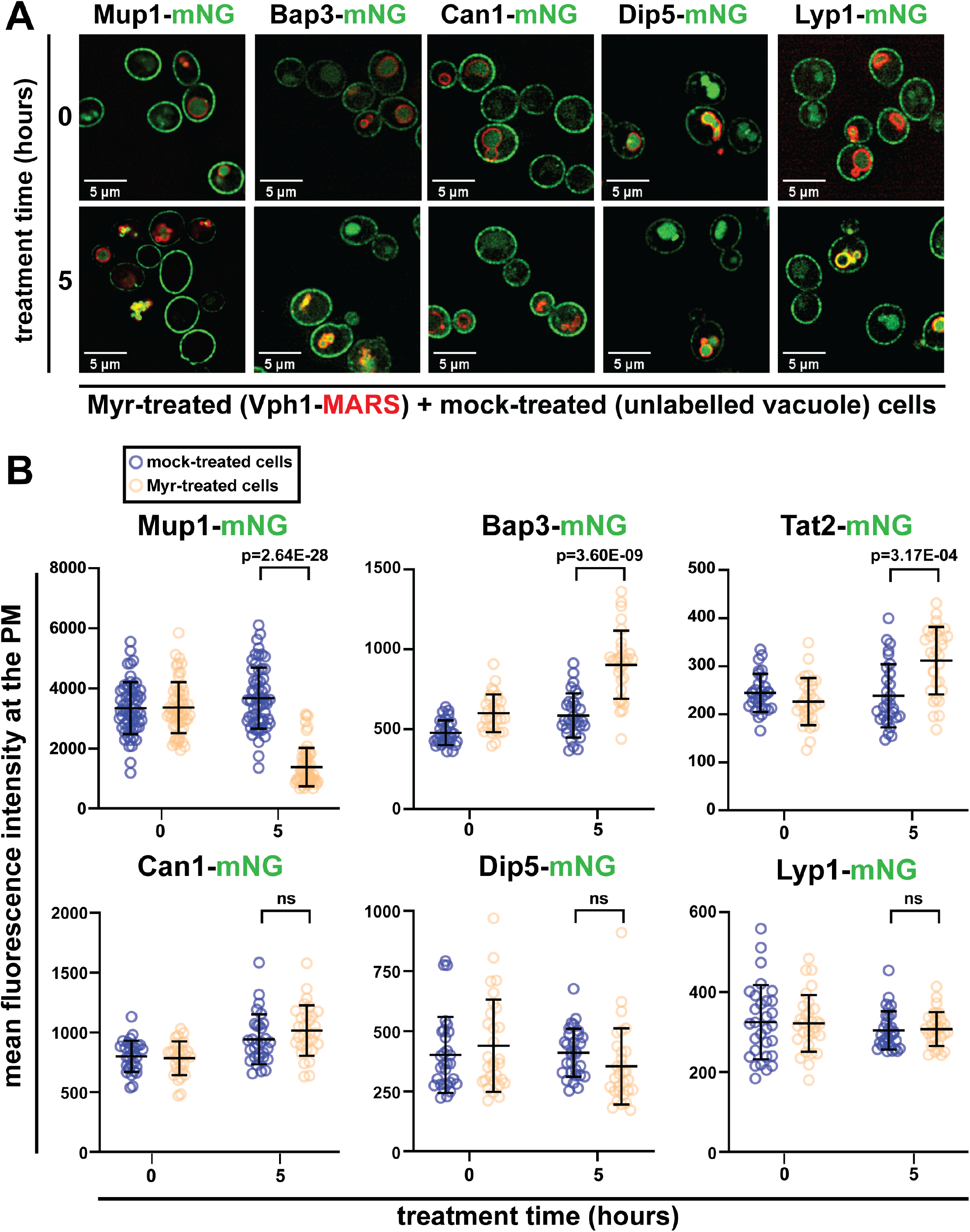
Myr treatment selectively decreases the level of the primary methionine permease, Mup1, at the plasma membrane. (A) Mixed population of yeast cells, expressing select mNG-tagged amino acid transporters (AATs), visualized under a fluorescence microscope after treatment with Myr (400 ng·mL^-1^) or mock solution (95% ethanol). Vph1 is a vacuolar marker used to distinguish the two yeast populations in a mixture visualized at 0 h and 5 h after treatment. (B) Quantification of the mean fluorescence intensity of select AAT-mNG at the PM, measured using Fiji (n = 30 - 60 cells; ±SD (error bars).

### Yeast cells respond to Myr by increasing abundance of specific glucose transporters

Given that yeast cells respond to Myr by altering the PM composition of AATs we considered the possibility that other nutrient transporters may also be affected. Specifically, given the relationship between glucose metabolism and aging, we hypothesized that Myr treatment may alter composition or activity of glucose transporters. To examine the effect of Myr on glucose transport, yeast cells were incubated with radiolabeled ^3^H-glucose and uptake was measured with or without Myr treatment. Notably, glucose uptake capacity was reduced in cells treated with Myr for at least 2 hours (**FIG 2A**). To determine if Myr treatment affects the abundance of glucose transporters at the PM, we examined cells harboring chromosomal mNeonGreen (mNG) fusions to a panel of four hexose transporter genes (*HXT1, HXT2, HXT3, HXT6*) in wildtype (SEY6210) and vacuolar Vph1-mCherry-expressing background strains. Cells were grown to mid-log phase, treated with Myr or mock-treated for 5 hours and mixed just prior to visualization by fluorescence microscopy (as described in **FIG 1**). We found that Myr treatment had no effect on the PM abundance of Hxt3 or Hxt6, while the low-affinity glucose transporter Hxt1 and the high-affinity glucose transporter Hxt2 exhibited increased abundance at the PM (**FIG 2B-C** and **FIG S2A-B**). Thus, Myr-treated cells selectively increase PM abundance of specific hexose transporters, while experiencing decreased capacity for glucose uptake. These results are unexpected and suggest impaired or suppressed activity of glucose transporters in sphingolipid-depleted cells.

**Figure 2.**
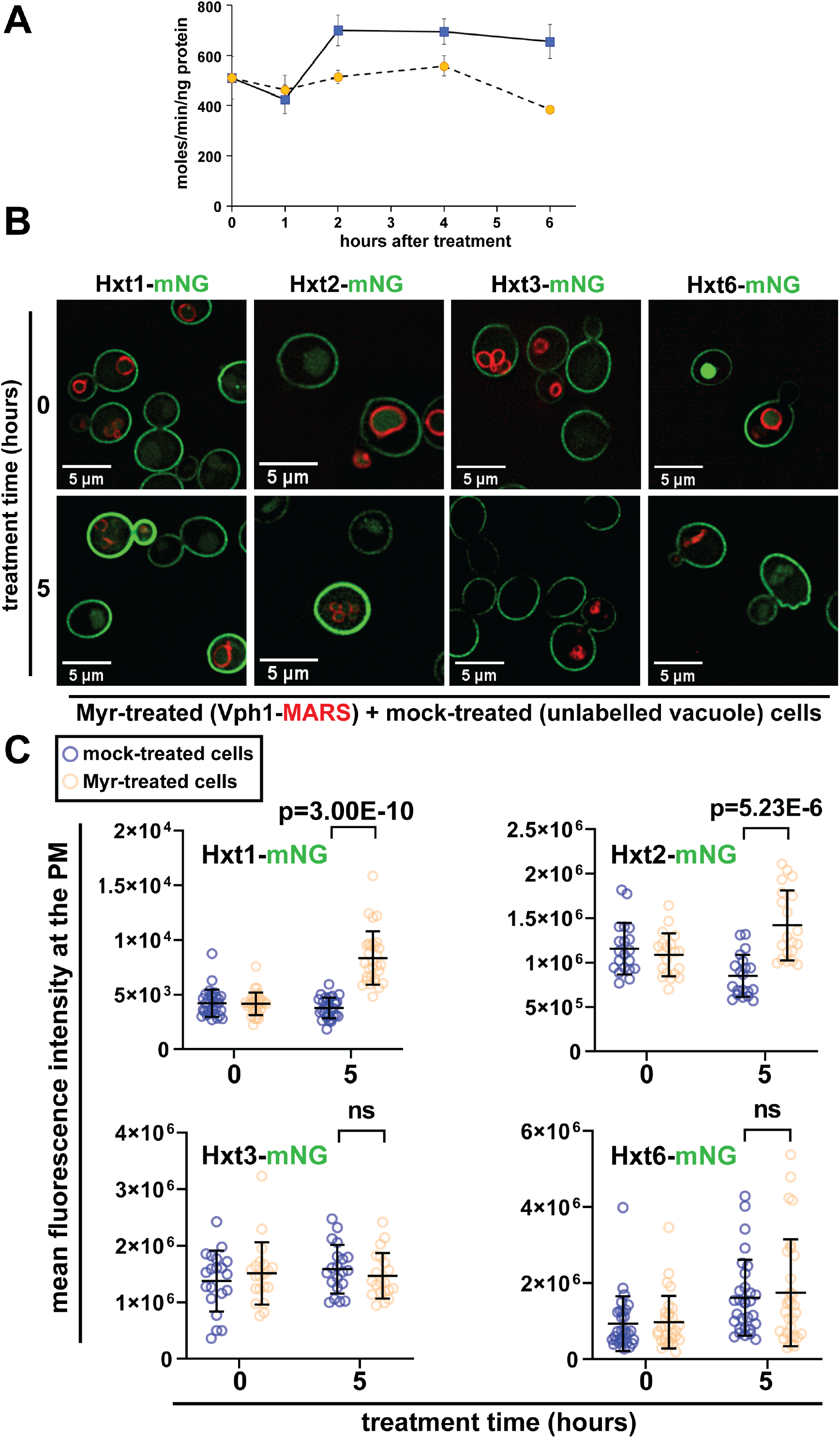
Myr decreases the cellular uptake of glucose despite the enhanced stability of hexose transporters Hxt1 and Hxt2 at the plasma membrane. (A) Time-course measurements of ^3^H-glucose uptake of cells treated with or without Myr. (B) Mixed population of yeast cells, expressing select mNG-tagged hexose transporters, visualized under a fluorescence microscope after treatment with Myr (400 ng·mL^-1^) or mock solution (95% ethanol). Vph1 is a vacuolar membrane marker used to distinguish the two yeast populations in a mixture visualized at 0 h and 5 h after treatment. (C) Quantification of the mean fluorescence intensity of select glucose transporters (C-term tagged with mNG) at the PM, measured using Fiji (n = 30 - 60 cells; ±SD (error bars).

We expanded our analysis to include other categories of integral membrane proteins at the PM. Myr-treatment resulted in a slight (but statistically insignificant) decrease in the PM abundance of Pma1, a P2-type ATPase that pumps protons out of the cell (**FIG S2C-D**). Myr treatment induced a correspondingly slight (but statistically significant) increase in the PM abundance of Pma2 (**FIG S2C-D**), a paralog of Pma1. While these changes were subtle, Myr treatment induced more substantial increases in the abundance of other proteins at the PM. For example, the pheromone receptor Ste2 exhibited largely vacuolar localization in mock-treated cells but localized to the PM in Myr-treated cells (**FIG S2C-D**) suggesting that Myr treatment interferes with constitutive endocytic trafficking of Ste2. Similar results were observed for the stress-sensing signal transducer Wsc1 (**FIG S2C-D**). Interestingly, Myr treatment also increased the PM abundance of the ABC family multidrug transporter Pdr5 (**FIG S2C-D**). A summary of the Myr treatment response of all integral PM proteins examined is provided in **Table 1**.

**Table 1.** Effect of myriocin on PM localization of nutrient transporters and other PM proteins. This table summarizes the microscopy-based localization data presented in Figure 1 and Figure 2 (and associated supplemental data). In cases where Myr treatment resulted in a significant change (P-value < 0.01 using Student’s T-test when compared to mock-treated cells) the box corresponding to the magnitude change (% Effect) is shaded red or green to indicate the extent of decrease or increase (respectively) observed upon Myr treatment.

| Functional Classification | PM Protein | Description | Response to Myriocin |  |
| --- | --- | --- | --- | --- |
|  |  |  | % Effect | P-value |
| amino acid transporters | Mup1 | transporter: Met | -62.44% | 2.64E-28 |
|  | Can1 | transporter: Arg | 7.93% | 0.0761 |
|  | Dip5 | transporter: Asp, Glu, Gln, Asn, Ser, Gly, Ala | -13.87% | 0.0558 |
|  | Lyp1 | transporter: Lys | 1.07% | 0.3917 |
|  | Bap3 | transporter: Cys, Leu, Ile and Val | 54.15% | 3.60E-09 |
|  | Tat2 | transporter: Trp and Tyr | 30.67% | 3.17E-04 |
|  | Bap2 | transporter: Leu | 9.68% | 0.0575 |
|  | Gnp1 | transporter: Ser, Leu, Thr, Cys, Met, Asn, Gln | -6.35% | 0.1868 |
|  | Hip1 | transporter: His | -12.90% | 0.0455 |
|  | Gap1 | general amino acid transporter | n.d. (mostly vacuolar) |  |
| hexose transporters | Hxt1 | glucose transporter (low affinity) | 120.78% | 3.00E-10 |
|  | Hxt2 | glucose transporter (high affinity) | 67.06% | 5.23E-06 |
|  | Hxt3 | glucose transporter (low affinity) | -0.07% | 0.2112 |
|  | Hxt6 | glucose transporter (high affinity) | 7.97% | 0.3144 |
| proton pumps | Pma1 | proton pump | -19.40% | 0.0546 |
|  | Pma2 | proton pump | 24.31% | 0.0095 |
| other PM proteins | Ste2 | receptor for alpha-factor pheromone | 81.35% | 4.33E-11 |
|  | Wsc1 | stress signaling transmembrane protein | 217.11% | 1.48E-12 |
|  | Pdr5 | ABC family multidrug transporter | 46.74% | 5.80E-06 |

### Myriocin decreases the bulk inflow of the PM

Since several PM proteins accumulated at the PM following Myr treatment we hypothesized this may be due to a decrease in bulk endocytosis. To measure bulk endocytosis, the lipophilic tracer dye FM4-64 (N-(3-triethylammoniumpropyl)-4-(p-diethylaminophenylhexatrienyl) pyridium dibromide) was added to pulse-label the PM, followed by washing and chasing for 1 hour. Bulk endocytosis results in most FM4-64 being delivered to the limiting membrane of the vacuole (labelled by Vph1-mNG) in mock-treated cells (**FIG S3A-B**). In contrast, cells treated with Myr for 4 hours exhibited reduced colocalization of FM4-64 with Vph1-mNG (**FIG 3A-B** and **FIG S3C-D**), indicating a significant reduction in bulk endocytosis. Notably, treatment of cells with Myr for 1 hour had no effect on bulk endocytosis, while Myr treatment for 2 hours resulted in a partial but significant defect in bulk endocytosis (**FIG 3A-B**). This defect in bulk endocytosis after 4 hours of Myr treatment likely contributes to the accumulation of various PM proteins in the PM (**Table 1**). Given that Myr treatment impairs bulk endocytosis, it is striking that the methionine transporter Mup1 undergoes selective endocytic clearance on the same time scale (**FIG 1** and **Table 1**). We set out to understand the mechanistic basis for selective endocytosis of Mup1 in Myr-treated cells.

**Figure 3.**
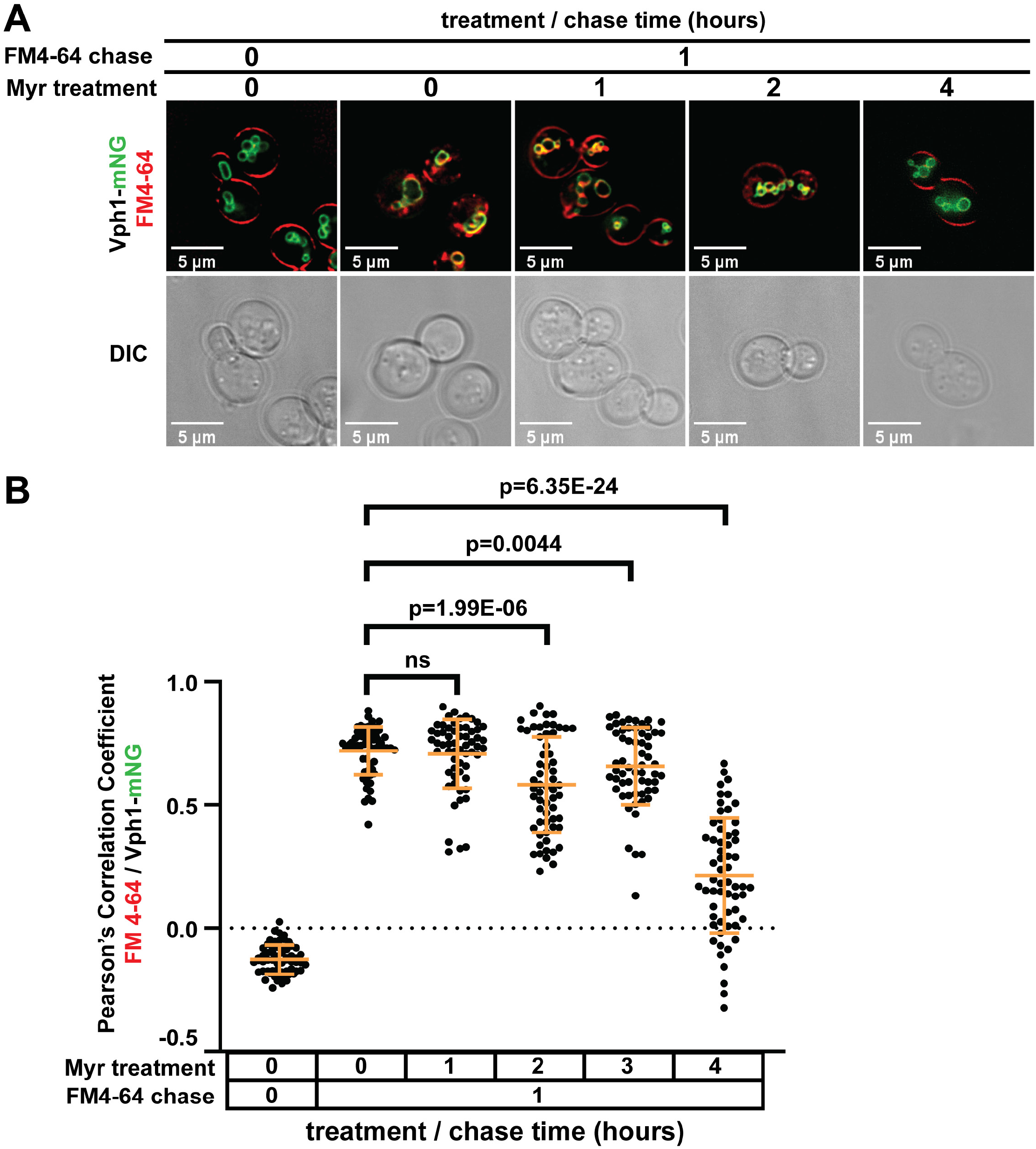
Myr inhibits the bulk endocytic trafficking of plasma membrane lipids. (A) Fluorescence microscopy of cells showing bulk endocytic trafficking of PM lipids, which eventually fuse with the vacuolar membrane. PM lipids were stained with a red lipophilic dye, FM 4-64 (fluorescent red), and vacuolar membrane was marked with Vph1-mNG (fluorescent green). Cells were treated with 400 ng·mL^-1^ Myr (for 0, 1, 2 or 4 hours), and the trafficking and vacuolar fusion of PM-derived lipids were determined after 1 hour of incubation with FM 4-64 in YPD media at 30°C. (B) Co-localization of FM 4-64 and Vph1-mNG measured as Pearson’s Correlation Coefficient values using softWoRx (ver. 7.0.0).

### Myriocin-induced trafficking of Mup1 requires the Rsp5 adaptor Art2

To better understand how Myr triggers the selective endocytic clearance of Mup1, we first attempted to validate our findings using yeast cells with chromosomal *MUP1* fused at its C-terminus to superecliptic pHluorin, a GFP variant that does not fluoresce in the acidic environment of the vacuole [23]. In this yeast variant, steady state levels of Mup1 at the PM can be measured by flow cytometry [24]. Similar to our fluorescence microscopy results, this approach revealed Mup1 levels at the PM begin to decrease after 4 hours of Myr treatment (**FIG 4A**). To test whether this response is due to sphingolipid depletion, we cultured yeast cells in media supplemented with phytosphingosine (PHS), providing an intermediate product of the SL biosynthesis pathway which effectively bypasses the Myr-imposed enzymatic block. Notably, PHS supplementation prevented the Myr-induced endocytosis of Mup1 (**FIG 4B**), indicating this response is the result of sphingolipid depletion. Similar to Myr, aureobasidin (AbA), which inhibits the inositol phosphorylceramide synthase AUR1, also triggered the clearance of Mup1, an effect which could not be reversed by PHS supplementation (**FIG 4B**). Since PHS supplementation does not bypass the AbA enzymatic block, these results reveal that PM clearance of Mup1 is triggered by depletion of complex SLs which are synthesized downstream of ceramide (e.g., inositol phosphorylceramides and their mannosylated derivatives).

**Figure 4.**
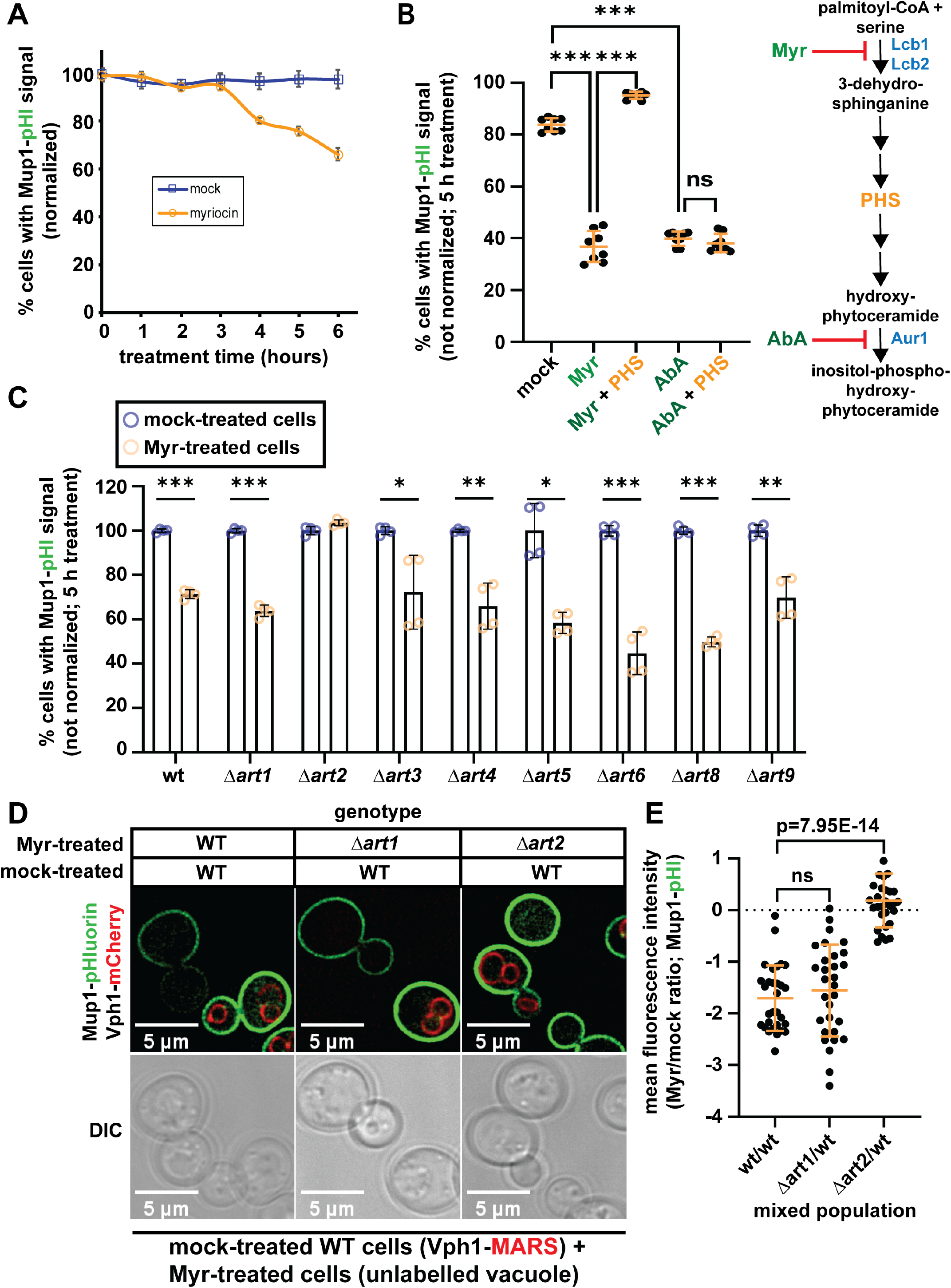
Sphingolipid depletion induces the endocytic trafficking of Mup1, requiring the arrestin adaptor protein Art2. (A) Mup1-pHluorin fluorescence decreasing at the PM after 4 h of Myr treatment as measured using a flow cytometer (n = 10,000 cells; ±SD (error bars)). (B) Phytosphingosine supplementation stabilizes Mup1-pHluorin at the PM of Myr-treated cells but not on Aureobasidin A (AbA)-treated cells. Mup1-pHluorin fluorescence signal at the PM of cells was measured by a flow cytometer after 5 hours of treatment (eight biological replicates; n = 10,000 cells; ±SD (error bars)). Myr and AbA inhibit the serine palmitoyl transferase (Lcb1/Lcb2 complex) and phosphatidylinositol:ceramide phosphoinositol transferase (Aur1), respectively, of the sphingolipid biosynthesis pathway. (C) *ART2* is required in Myr-induced endocytic trafficking of Mup1-pHluorin. Mup1-pHluorin fluorescence signal at the PM of cells was measured by a flow cytometer after 5 hours of treatment (four biological replicates; n = 10,000 cells; ±SD (error bars)). (D) Fluorescence microscopy showing that the *Δart2* null-deletion strain has stable Mup1-pHluorin signal at the PM after Myr treatment. (E) Quantification Mup1-pHluorin fluorescence at the PM of cells shown in Fig. 4D.

The endocytosis of nutrient transporters in yeast is controlled primarily by Rsp5-mediated ubiquitylation events (reviewed in [25]). Pairing of the E3 ubiquitin ligase Rsp5 with cargo adaptors called ARTs (*<u>a</u>*rrestin-*<u>r</u>*elated *<u>t</u>*rafficking adaptors) determines the specificity of substrate targeting and promotes adaptation to nutrient fluctuations and stress conditions [26, 27]. Analysis of the yeast transcriptional response to Myr treatment [28] revealed increased transcript abundance for Rsp5 and several ARTs (**FIG S4A-B**), indicating that sphingolipid depletion alters gene expression in a way that could affect endocytic trafficking. To determine if these transcript-level changes underlie alterations at the protein level, we quantified the abundance of mNG C-terminal fusions to various ART proteins (expressed from endogenous chromosomal loci) using total fluorescence measurements (**FIG S4C**). Myr-induced changes in transcript and protein levels correlated in many cases (e.g., ART1 and ART2) but not in every case (e.g., ART4). Notably, both *ART1* and *ART2* were upregulated in a Myr-treatment time course (**FIG S4A-C**) and both are known to regulate Mup1 endocytosis in response to excess methionine [27, 29] and nitrogen starvation [30], respectively. To test if ARTs are involved in this response, we characterized Myr-triggered Mup1-pHluorin clearance in a panel of ART deletion yeast strains and found that all ARTs tested were dispensable except for *ART2* (**FIG 4C).** To validate this result, we performed fluorescence microscopy to compare the PM abundance of Mup1-pHluorin in wildtype, *Δart1* or *Δart2* yeast cells. Strikingly, Mup1 clearance was not detected in Myr-treated *Δart2* yeast cells, while loss of ART1 did not affect this response (**FIG 4D-E**). These findings indicate that Art2 mediates the endocytic clearance of Mup1 in response to Myr treatment.

Previous work demonstrated that nitrogen starvation triggers Art2-dependent endocytosis of Mup1 which required transcriptional induction of Art2 by activation of the general amino acid starvation response [30]. This stress response requires activity of the upstream activating kinase Gcn2, which phosphorylates eIF2α to mediate the response. We hypothesized that Myr-triggered endocytosis of Mup1 may likewise occur through activation of the general amino acid starvation response and subsequent up-regulation of Art2. To test this, we compared Myr-triggered Mup1 endocytosis in wildtype and *Δgcn2* mutant cells. Unexpectedly, we found that Gcn2 is dispensable for Myr-triggered endocytosis of Mup1 (**FIG S5**). Thus, in contrast to Mup1 endocytosis that occurs during nitrogen starvation [30], Myr-induced endocytosis of Mup1 occurs independently of the general amino acid starvation response.

### Myr-induced trafficking of Mup1 requires C-terminal Lys63-linked polyubiquitylation

Ubiquitylation at N-terminal lysines (K27 and K28) is required for Mup1 endocytosis in response to excess methionine [31–33] while ubiquitylation at C-terminal lysines (K567 and K572) is reported to mediate endocytic clearance in response to nitrogen starvation [30]. Structure predictions from the AlphaFold protein structure database indicate that the N-terminal ubiquitylation sites (K27 and K28) exist in a largely unstructured region, while the C-terminal ubiquitylation sites (K567 and K572) occur in an alpha-helical region (**FIG 5A**). Since Art1-mediated ubiquitylation of Mup1 occurs at N-terminal lysines (K27 and K28) and Art2 was previously reported to bind at the C-terminus of Mup1 [30] we predicted that ubiquitylation of C-terminal lysine residues may be required for Myr-induced endocytosis of Mup1. To test this prediction, we characterized Myr-induced trafficking of Mup1-mNG in strains with short C-terminal truncations lacking one or both C-terminal lysine residues (ΔK572 and ΔK567, respectively) (**FIG 5B**). While Mup1^ΔK572^-mNG exhibited Myr-induced endocytic clearance, Mup1^ΔK567^-mNG was unresponsive to Myr treatment (**FIG 5C-D**). These results suggest that Myr-triggered Mup1 endocytosis requires ubiquitylation at its C-terminal lysine residues.

**Figure 5.**
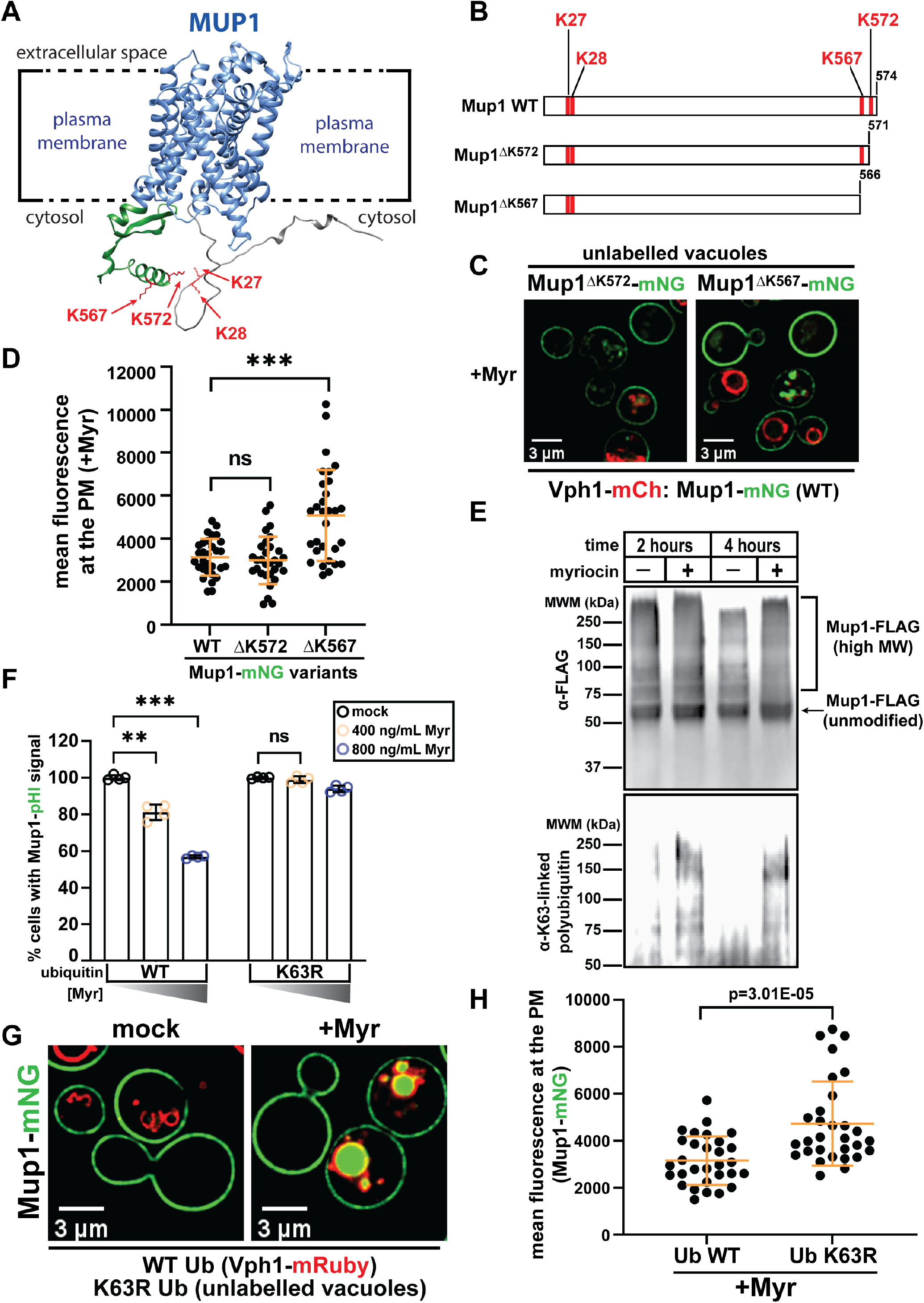
Myr-induced trafficking of Mup1 requires K63-linked polyubiquitylation at the C-terminal lysines of Mup1. (A) Diagram showing the lysine residues at the cytosolic region of PM-associated Mup1. (B) Diagram showing the C-terminal truncation variants of Mup1. (C) Fluorescence microscopy of mixed cell populations (mNG-tagged Mup1 wt and truncation variants) after 5 h of Myr treatment. (D) Quantification of Mup1-mNG expressed as mean fluorescence intensity at the PM of cells visualized in Fig. 5C. (E) Anti-FLAG and anti-K63 polyUb immunoblots of Mup1-FLAG enriched by anti-FLAG immunoprecipitation after 2 or 4 h of Myr treatment. (F) Flow cytometry of SUB280-derived cells (all four ubiquitin-coding genes deleted) that *trans-express* only the wt or K63R ubiquitin, and endogenously express Mup1-pHluorin after 5 hours treatment with or without Myr (four biological replicates; n = 10,000 cells; ±SD (error bars)). (G) Fluorescence microscopy showing that SUB280 cells expressing only the Ub K63R have more stable Mup1-mNG signal at the PM after Myr treatment. (H) Quantification of Mup1-mNG mean fluorescence intensity at the PM of cells shown in Fig. 5G.

To measure the ubiquitylation of Mup1, we affinity purified Mup1-FLAG from yeast lysates and analyzed it by SDS-PAGE and quantitative immunoblotting. This analysis revealed a significant increase in K63-linked ubiquitin polymers associated with Mup1 in response to Myr treatment (**FIG 5E**). Previous work demonstrated that conjugation to monoubiquitin is sufficient for Mup1 endocytosis in response to excess methionine [34]. In contrast, we found that yeast cells expressing Ub^K63R^ as the sole source of ubiquitin were deficient for Myr-triggered endocytosis of Mup1 (**FIG 5F-H**), indicating that K63-linked ubiquitin polymers are required for this response. Taken together, our results reveal that Myr-induced endocytosis of Mup1 is ubiquitin-mediated but proceeds by a mechanism that is distinct from methionine-induced endocytosis.

### Ede1 and Ent1 function redundantly in Myr-mediated Mup1 endocytic clearance

Multiple ubiquitin-binding proteins in yeast, including the epsins Ent1 and Ent2 and the epsin-like Ede1, function as adaptors that capture ubiquitylated cargoes during endocytic vesicle formation. Among these, Ede1 was previously found to be crucial in mediating Mup1 trafficking in response to excess methionine [35]. To examine a role for endocytic adaptors in the cellular response to Myr we analyzed the abundance of mNG C-terminal fusions to endocytic adaptor proteins (expressed from endogenous chromosomal loci) using total fluorescence measurements. This analysis revealed significant Myr-triggered increases in protein levels of Ent1 and Ede1 while no significant change in the level of Ent2 was detected (**FIG 6A**). We next measured co-localization between endocytic adaptors and Mup1 in response to Myr treatment. This analysis revealed that Myr treatment induced co-localization of Mup1 with Ent1 and Ede1, but not Ent2 (**FIG 6B**). Importantly, we did not observe significant changes for Ede1 co-localization with either Can1 (an arginine transporter) or Pil1 (an eisosome component) during the same time course (**FIG S6B**). To determine if the increased association between Mup1 and Ede1 was due to ubiquitin binding, we analyzed association between Mup1-mNG and a variant of Ede1 lacking its C-terminal UBA domain, which is known to preferentially interact with K63-linked polymers [36]. Notably, the Ede1^Δuba^ variant did not exhibit increased association with Mup1 in response to Myr treatment (**FIG 6C**). Finally, we analyzed Myr-triggered Mup1 trafficking in yeast strains lacking endocytic adaptors. This analysis revealed that Ent1, Ent2 and Ede1 are all individually dispensable for Myr-induced Mup1 trafficking, despite the fact that loss of Ede1 is sufficient to prevent methionine-triggered Mup1 endocytosis (**FIG S6C** and [35]). Analysis of double mutants revealed that Myr-triggered Mup1 endocytosis occurs normally in *Δent2Δede1* cells but is blocked in *Δent1Δede1* cells (**FIG 6D-F**). (Notably, *Δent1Δent2* double mutant yeast cells are inviable [35] and thus could not be tested in our analysis.) These data reveal that Ent1 and Ede1 contribute redundantly to Mup1 clearance in response to Myr treatment.

**Figure 6.**
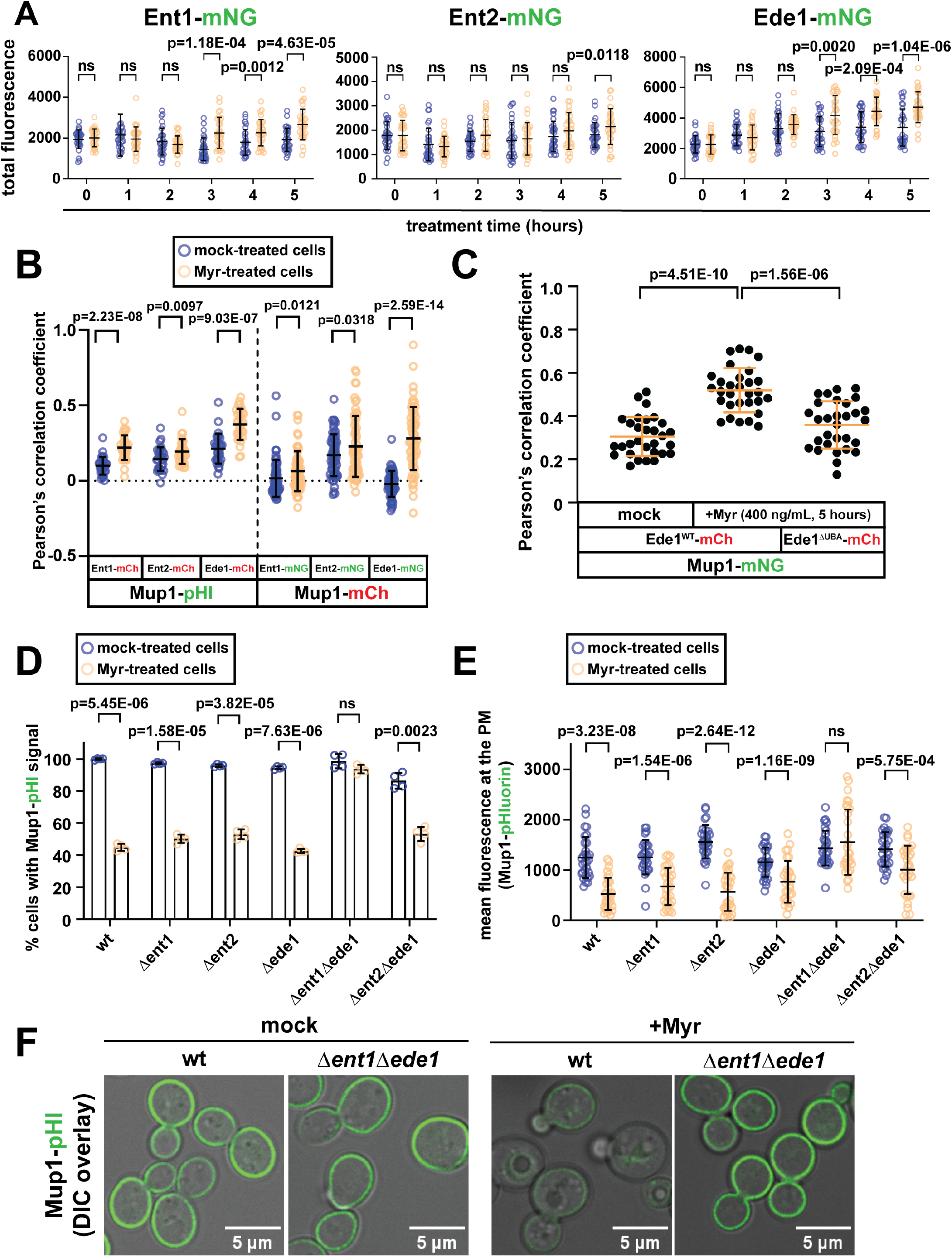
Myr-induced trafficking of Mup1 requires the ubiquitin-binding endocytic adaptor proteins Ent1 and Ede1. (A) Myr increases the fluorescence level of Ent1-, Ent2-, and Ede1-mNG in cells. Total fluorescence intensity of Ent1-, Ent2-, or Ede1-mNG was measured using Fiji after 0 to 5 hours of treatment (n = 30 cells; ±SD (error bars)). (B) Myr increases the co-localization of Mup1-pHluorin with Ent1-, Ent2-, and Ede1-mCherry, or Mup1-mCherry with Ent1-, Ent2-, and Ede1-mNG after 5 hours of Myr treatment. (C) The ubiquitin-binding UBA domain of Ede1 is required in the induction of Mup1-mNG/Ede1-mCherry co-localization in response to 5-hour Myr treatment. Co-localization was measured in 30 cells and graphically presented as Pearson’s Correlation Coefficient values. (D) Ent1 and Ede1 are required in Myr-induced trafficking of Mup1. The fluorescence of Mup1-pHluorin was measured in cells after 5 hours with or without Myr treatment (four biological replicates; n = 10,000 cells; ±SD (error bars)). (D) Quantification of Mup1-pHluorin mean fluorescence intensity at the PM of cells after 5 hours with or without Myr treatment (n = 30 cells; ±SD (error bars)). (E) Fluorescence microscopy of wt and *Δent1Δede1* cells expressing Mup1-pHluorin after five hours with or without Myr treatment.

## Discussion

It is well-established that genetic and pharmacological interventions that perturb sphingolipid biosynthesis promote longevity in a variety of model organisms [9], although how sphingolipid homeostasis and life span are coupled is not fully understood. Previously, we have reported that sphingolipid reduction in yeast extends chronological life span [37] and that this is associated with a state of amino acid restriction, which is accomplished at least in part by decreasing the uptake of extracellular amino acids [22]. Here, we report the unexpected result that, for almost all nutrient transporters and PM-associated proteins examined, Myr treatment either had on effect or increased abundance at the PM (**Table 1**). The only exception was Mup1, which undergoes endocytic clearance following 4 hours of Myr treatment (**FIG 4**). Thus, the observed decrease in abundance of most amino acids observed upon Myr treatment is not likely due to broad endocytic clearance of AATs, and in fact occurs despite the increased PM abundance of Bap3 and Tat2. Similar results were observed for hexose transporters, some of which accumulated at the PM despite decreased glucose uptake during a Myr treatment time course. One possible explanation for this apparent disparity is that sphingolipid depletion may lower transport activity without inducing endocytosis. In some cases, the accumulation of specific PM proteins following Myr treatment may be due to inhibition of bulk endocytosis, which was observed using FM4-64 trafficking assays (**FIG 3**). Alternatively, it is also possible that cells respond to decreased amino acid and glucose availability by upregulating the biosynthesis and secretion of specific nutrient transporters. Our analysis suggests that the activities of many nutrient transporters are coupled to sphingolipid abundance at the PM, and future studies will need to address mechanism of this coordination.

The methionine transporter Mup1 was unique amongst PM proteins in being selectively targeted for endocytic clearance following sphingolipid depletion. More specifically, our finding that both Myr and AbA triggered Mup1 endocytosis, but addition of PHS only suppressed the effect of Myr and not the effect of AbA, indicates that depletion of inositol phosphorylceramides (and/or downstream products) triggers Mup1 endocytosis. The mechanism of methionine-induced endocytosis of Mup1 is well-characterized [31, 34, 35, 38]: **(*i*)** it involves ubiquitylation of N-terminal lysine residues by the Art1-Rsp5 E3 ubiquitin ligase complex, **(*ii*)** it occurs independently of ubiquitin polymer formation, and **(*iii*)** it requires the endocytic adaptor Ede1. In contrast, we find that Myr-induced endocytosis of Mup1 **(*i*)** is mediated by the Art2-Rsp5 E3 ubiquitin ligase complex, **(*ii*)** requires C-terminal lysine residues, **(*iii*)** requires the formation of K63-linked ubiquitin polymers, and **(*iv*)** requires either Ede1 or Ent1 as an endocytic adaptor. Furthermore, Myr treatment induces co-localization of Mup1 with Ent1 and Ede1, and the Mup1-Ede1 co-localization requires its C-terminal UBA domain, which is known to interact with ubiquitin. Thus, the mechanism of Myr-triggered Mup1 endocytosis is mechanistically distinct from methionine-mediated Mup1 endocytosis (**Fig. 7**).

**Figure 7.**
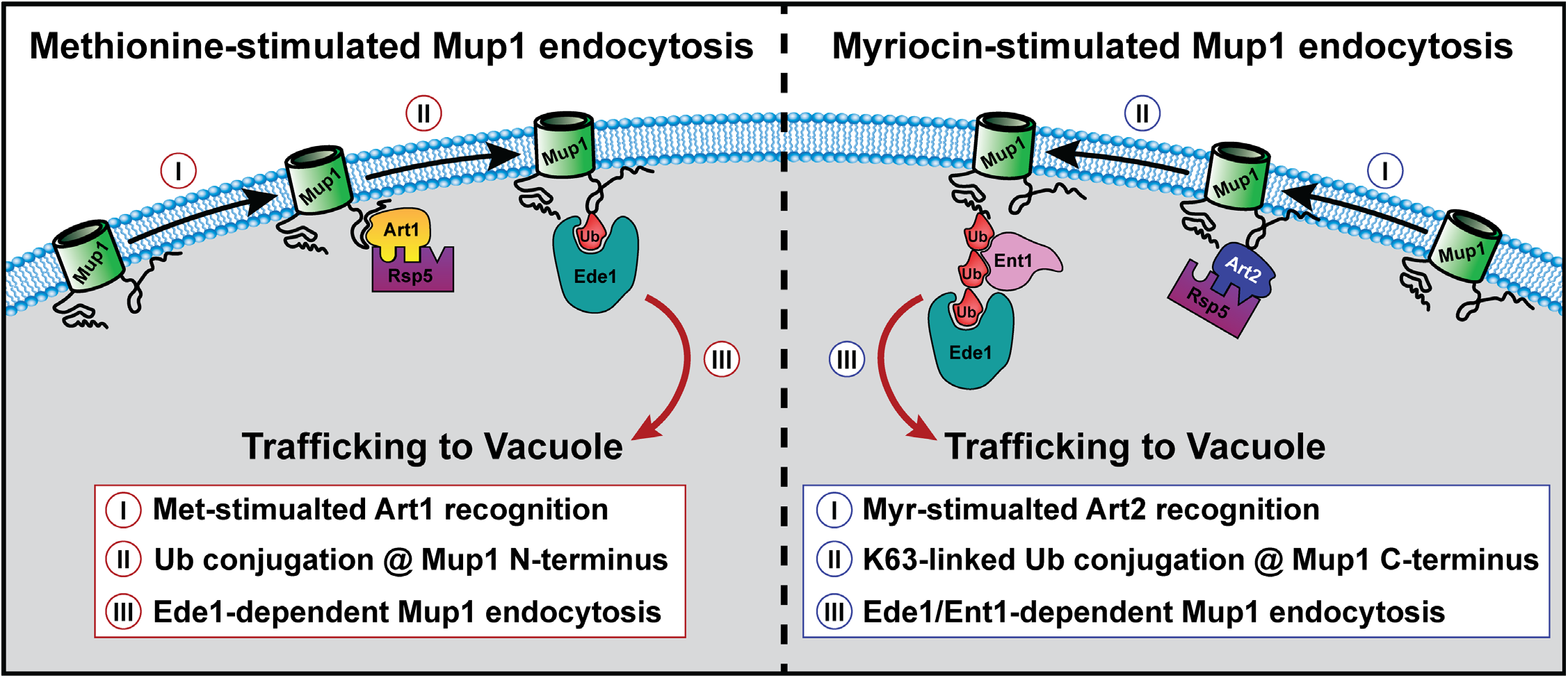
Model of Mup1 endocytic trafficking in response to excess methionine and sphingolipid depletion. The box insets highlight mechanistic distinctions between the two endocytic processes.

Importantly, Mup1 endocytosis has also been reported to occur during cellular adaptation to other stresses and environmental changes. Endocytosis of many PM proteins, including Mup1, occurs in response to depletion of nicotinic acid by a mechanism that relies on tetraspan Cos proteins but is distinct from known ART-Rsp5 complexes [39]. Another recent study reported that nitrogen starvation triggers endocytosis of multiple AATs – including Mup1, Can1, Lyp1, Tat2 – as well as glucose transporters Hxt1, Hxt2, and Hxt3 [30]. The nitrogen starvation-induced endocytosis of Mup1 and several other AATs (Can1, Lyp1, and Tat2) was Art2-dependent and required Gcn2-dependent induction of Art2 expression [30]. Although both nitrogen starvation and sphingolipid depletion induce Art2-dependent endocytosis of Mup1, there are two notable mechanistic distinctions. First, while nitrogen starvation induces Art2-mediated endocytosis of multiple nutrient transporters, the endocytosis induced by sphingolipid depletion is very selective for Mup1. Indeed, some nutrient transporters like Tat2, Hxt1 and Hxt2 were internalized during nitrogen starvation but accumulated at the PM in response to sphingolipid depletion. Second, while Gcn2 is required for Mup1 endocytosis during nitrogen starvation [30] it is dispensable for Myr-triggered Mup1 endocytosis (**FIG S5**). These distinctions indicate that Art2 is broadly activated in a Gcn2-dependent manner in response to nitrogen starvation, but that its Gcn2-independent activation during sphingolipid depletion is restricted to Mup1. Together, these results reveal distinct PM remodeling processes that occur during cellular adaptation to nitrogen starvation or to sphingolipid depletion.

It remains unclear why the methionine transporter Mup1 is selectively targeted for endocytic clearance following sphingolipid depletion, and we hypothesize that altered methionine homeostasis may be critical for Myr-mediated longevity. In support of this hypothesis, we recently reported that artificial stabilization of Mup1 at the PM suppresses the longevity-enhancing effects of sphingolipid depletion [28]. This is consistent with a recent study which reported that decreased intracellular methionine concentration mediates life span extension associated with caloric restriction [40]. Collectively, these studies underscore the critical importance of methionine metabolism as a determinant of aging, and they suggest commonalities between life span extension associated with caloric restriction and sphingolipid depletion. Ultimately, improved understanding of cellular adaptation to sphingolipid depletion, particularly with respect to PM transport functions that regulate intracellular nutrient concentrations, will reveal how compounds like myriocin promote health and longevity.

## Materials and Methods

### Strains, media and growth conditions

*Saccharomyces cerevisiae* strains expressing endogenous reporter proteins fused with fluorescent proteins were generated by homologous recombination or mating. Cells in synthetic complete dextrose (SCD) media (preparation described in Hepowit et al. 2022) were grown at 26°C with agitation (220 rpm) to mid-log phase (OD600 = 0.3 – 0.6) and treated with 400 ng·mL^-1^ myriocin (Cayman Chemical Company), Aureobasidin A (TaKaRa), phytosphingosine (Tokyo Chemical Industry), or mock solution (95% ethanol) as needed. Cells expressing Hip1-mNG were cultured in low-histidine SCD (2 μg·mL^-1^), while strains expressing Hxt6-mNG and Hxt7-mNG were grown in low-glucose SCD (0.2% glucose).

### Fluorescence Microscopy

Yeast cells endogenously expressing fluorescent fusion proteins (mNG, MARS or mCherry) were grown to mid-log phase in indicated SCD broth, treated with Myr for 5 h, concentrated by centrifugation (3,500 X *g* for 10 s), and visualized using a DeltaVision Elite Imaging system (Olympus IX-71 inverted microscope; Olympus 100× oil objective (1.4 NA); DV Elite sCMOS camera, GE Healthcare). For the bulk endocytosis experiments using the 4-[6-[4-(diethylamino)phenyl]-1,3,5-hexatrien-1-yl]-1-[3-(triethylammonio)propyl]-pyridiniumbromide dye (FM 4-64, Invitrogen), the cells were prepared as previously described [41]. Fluorescence colocalization was measured using Pearson correlation coefficients analyzed by the Softworx software (GE Healthcare). Images obtained from the red and green filter channels were merged and the background-subtracted mean fluorescence at the plasma membrane was measured using Fiji [42].

### Flow Cytometry

The endocytic trafficking of Mup1-pHluorin was analyzed using a flow cytometer as previously described [22, 43] with minor modifications. Briefly, cells endogenously expressing Mup1-pHluorin were grown to mid-log phase, treated with Myr or AbA, or in combination with PHS for 5 h, and analyzed using the BD Accuri™ C6 Plus Flow Cytometer. The flow was set in fast fluidics and the relative intensity of Mup1-pHluorin in 10,000 cells was measured in the FITC channel using 90% histogram gating of the mock-treated cells for signal normalization.

### Western Blotting

Endogenously expressed Mup1-FLAG was isolated by anti-FLAG immunoprecipitation as previously described [44], dissolved in urea sample buffer containing 10% β-mercaptoethanol, and resolved in 12% Bis-Tris PAGE gel by electrophoresis. Proteins were transferred onto PVDF membrane (0.45 μm, GE Healthcare Amersham) by electrophoretic transblotting, blocked with 3% bovine serum albumin, and incubated with the following primary antibodies: anti-FLAG (1:1,000; Sigma; Mab), anti-ubiquitin (1:10,000; LifeSensors; Mab; clone VU-1), anti-K63 (1:1,000; EMD Millipore, RAb; clone apu3). Secondary antibodies used were anti-mouse (IRDye 680RD-goat anti-mouse; LI-COR) and anti-rabbit (IRDye 800CW-goat anti-rabbit; LI-COR). Fluorescence of blots was visualized using the Odyssey CLx Imaging System (LI-COR) and quantified using Image Studio Lite (LI-COR).

### RNA seq analysis

Total RNA was isolated from the lysate of 5 OD600 unit of cells, sampled every 1 h increment of Myr treatment (up to 6 h), using the RNAeasy Mini Kit (Qiagen, Maryland). RNA samples were frozen in dry-ice ethanol bath and stored at −80°C until use. RNA seq was performed at the J. Carver Biotechnology Center at the University of Illinois and data are deposited in the Gene Expression Omnibus (GSE199904; NCBI tracking system #22817261).

## Supplementary Figure Legends

**Figure S1.**
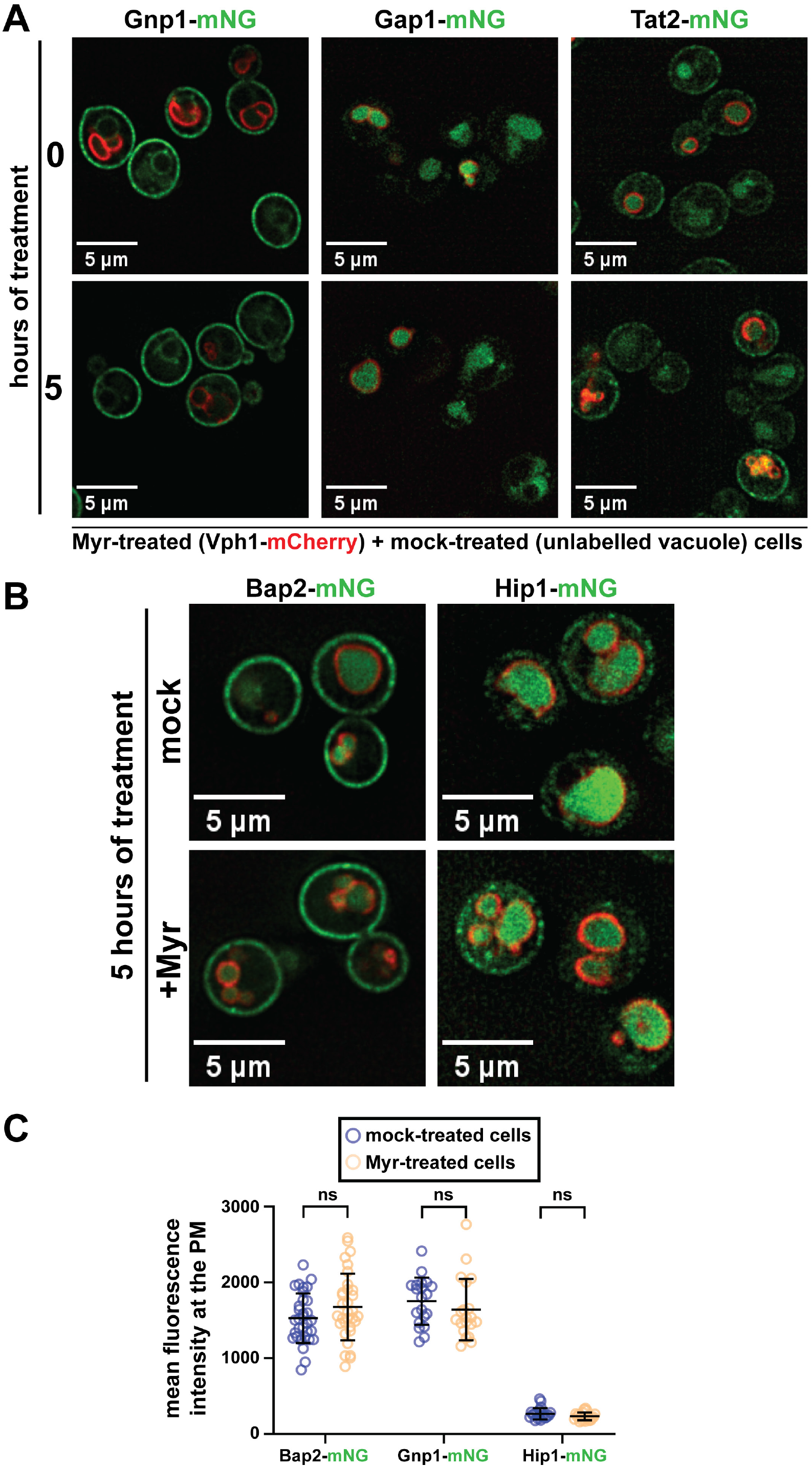
Microscopy of cells expressing endogenous mNG-tagged amino acid transporters (AATs) treated with or without Myr. (A) Yeast cells expressing AAT Gnp1, Gap1 or Tat2 in a mixed population assay. Myr-treated cells express Vph1-mCherry as a vacuolar marker to distinguish from mock-treated unlabeled cells. (B) Yeast cells expressing AAT Bap2 or Hip1 after 5 hours of treatment with or without Myr. Vph1-mCherry is used as a vacuolar marker. (C) Mean fluorescence intensity of AATs at the PM measured on cells in Suppl. Fig 1A-B (n = 20 cells; ±SD (error bars)).

**Figure S2.**
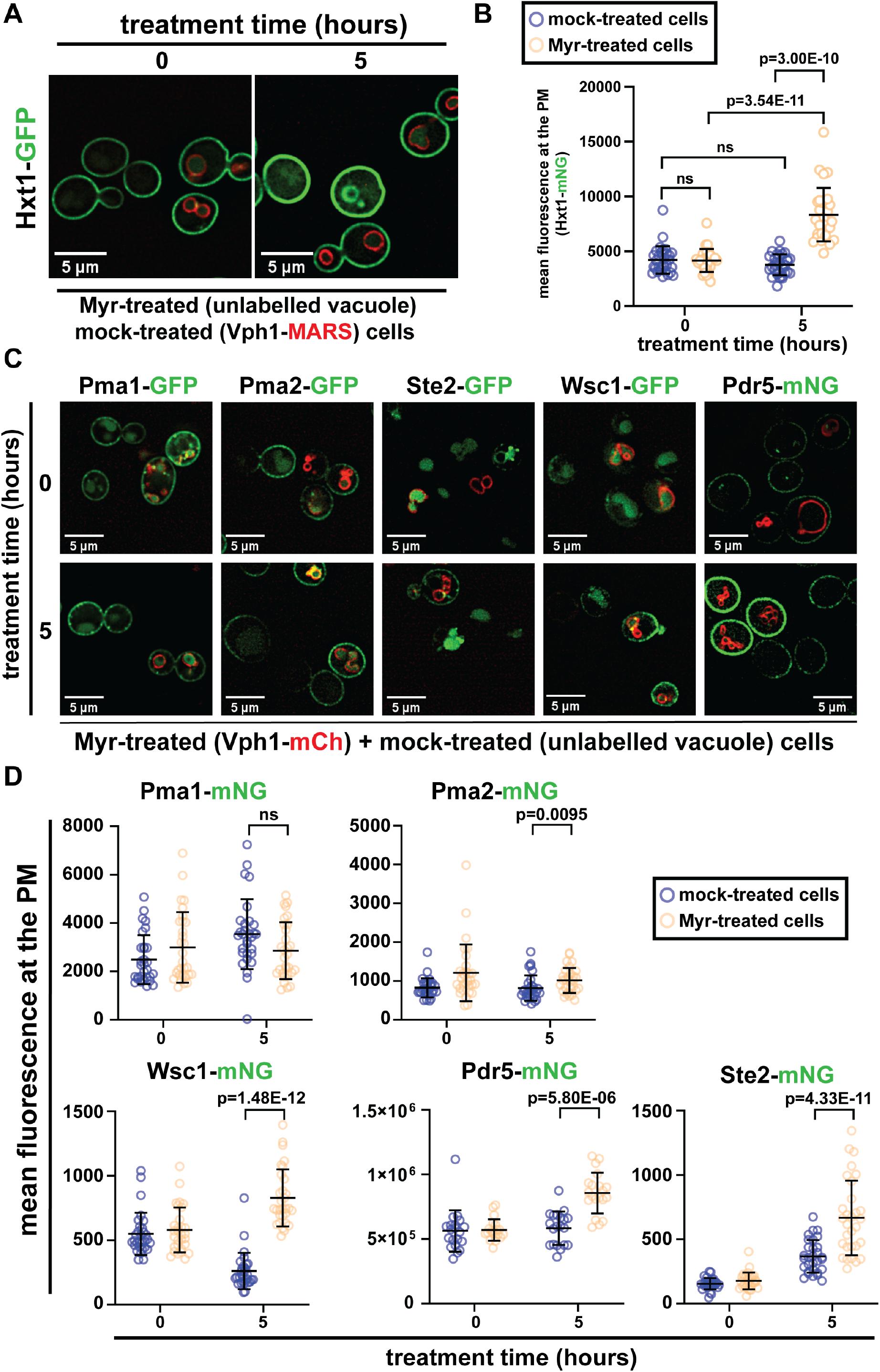
Myr stabilizes a subset of integral proteins at the PM. (A) Yeast cells expressing endogenous Hxt1-GFP treated with or without Myr in a mixed population microscopy assay. Mock-treated cells express Vph1-MARS as a red marker of vacuolar membranes, while Myr-treated cells have unlabeled vacuole. (B) Mean fluorescence intensity of Hxt1-GFP at the PM of cells shown in Suppl. Fig. 1A (n = 20 cells; ±SD (error bars)). (C) Fluorescence microscopy of mixed population of cells expressing select mNG-tagged proteins in trans. Myr-treated cells express Vph1-mCherry as a vacuolar marker to distinguish from mock-treated unlabeled cells. (D) Mean fluorescence intensity of mNG signal at the PM of cells shown in Suppl. Fig. 1C (n = 20-30 cells; ±SD (error bars)).

**Figure S3.**
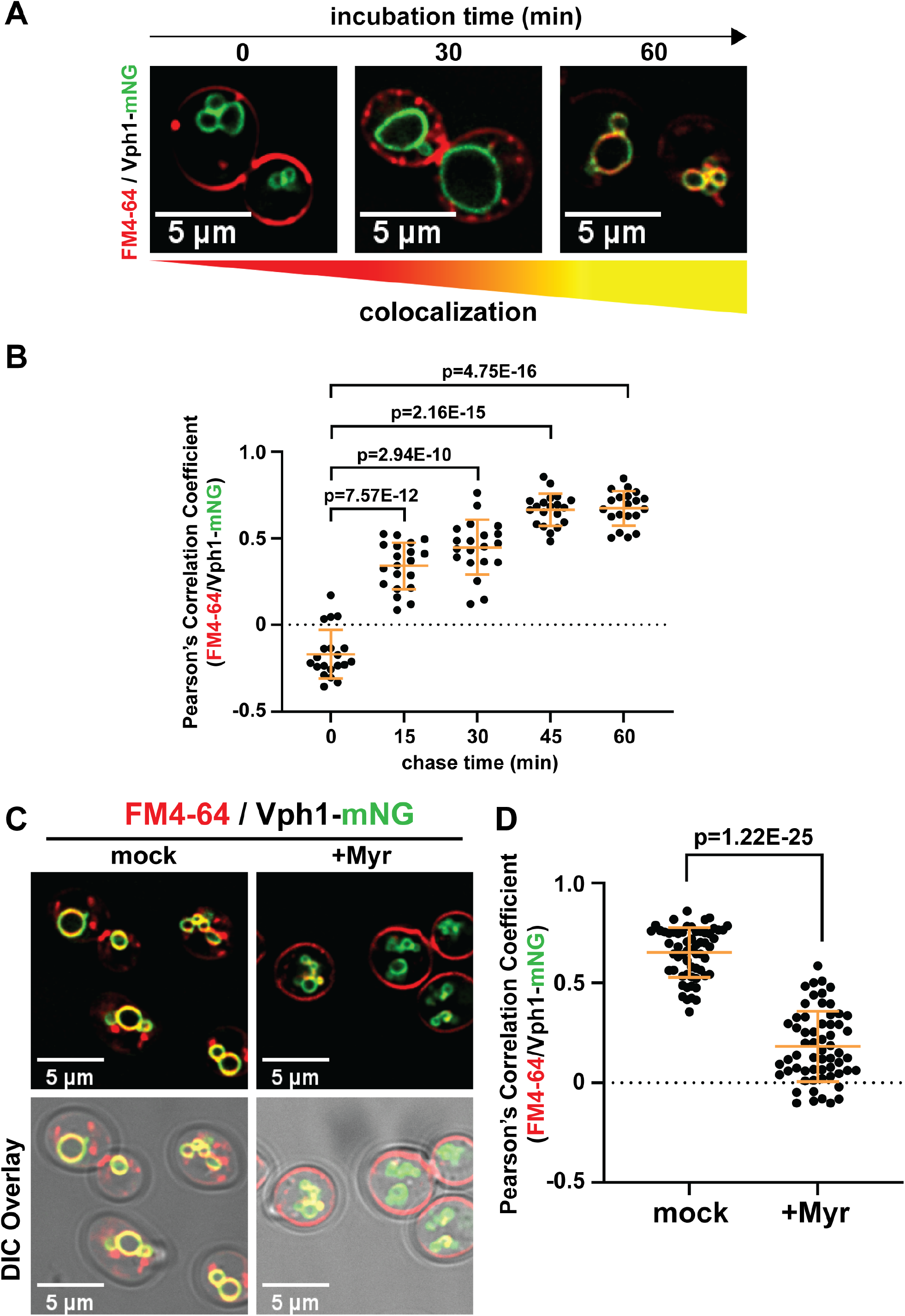
Myr inhibits the trafficking and vacuolar fusion of PM-derived lipids. (A) Endocytic trafficking and vacuolar fusion of PM lipids in S. cerevisiae SEY6210 cells. PM is dyed with lipophilic FM4-64 (fluorescent red) and vacuolar membrane is marked with Vph1-mNG (fluorescent green). Yellow coloration indicates the colocalization of FM4-64 and Vph1-mNG at the vacuole. After 1 h of Myr treatment, cells were washed and resuspended in fresh SCD media and the bulk PM lipid trafficking was visualized after 0, 30, 60 min of incubation at 26°C. (B) Quantification of Fm4-64/Vph1-mNG colocalization after treatment (0, 15, 30, 45 or 60 min) with FM4-64. Colocalization expressed as Pearson’s Correlation Coefficient values (n = 20 cells; ±SD (error bars)) measured using softWorx (ver. 7.0.0). (C) Fluorescence microscopy of Myr-treated S. cerevisiae BY4741 cells (4 hours) after 1 our incubation with FM4-64. (D) Quantification of Fm4-64/Vph1-mNG colocalization on Myr-treated cells (Suppl. Fig. 3C) after incubation with FM4-64 for 1 hour. Colocalization expressed as Pearson’s Correlation Coefficient values (n = 30 cells; ±SD (error bars)) measured using softWorx (ver. 7.0.0).

**Figure S4.**
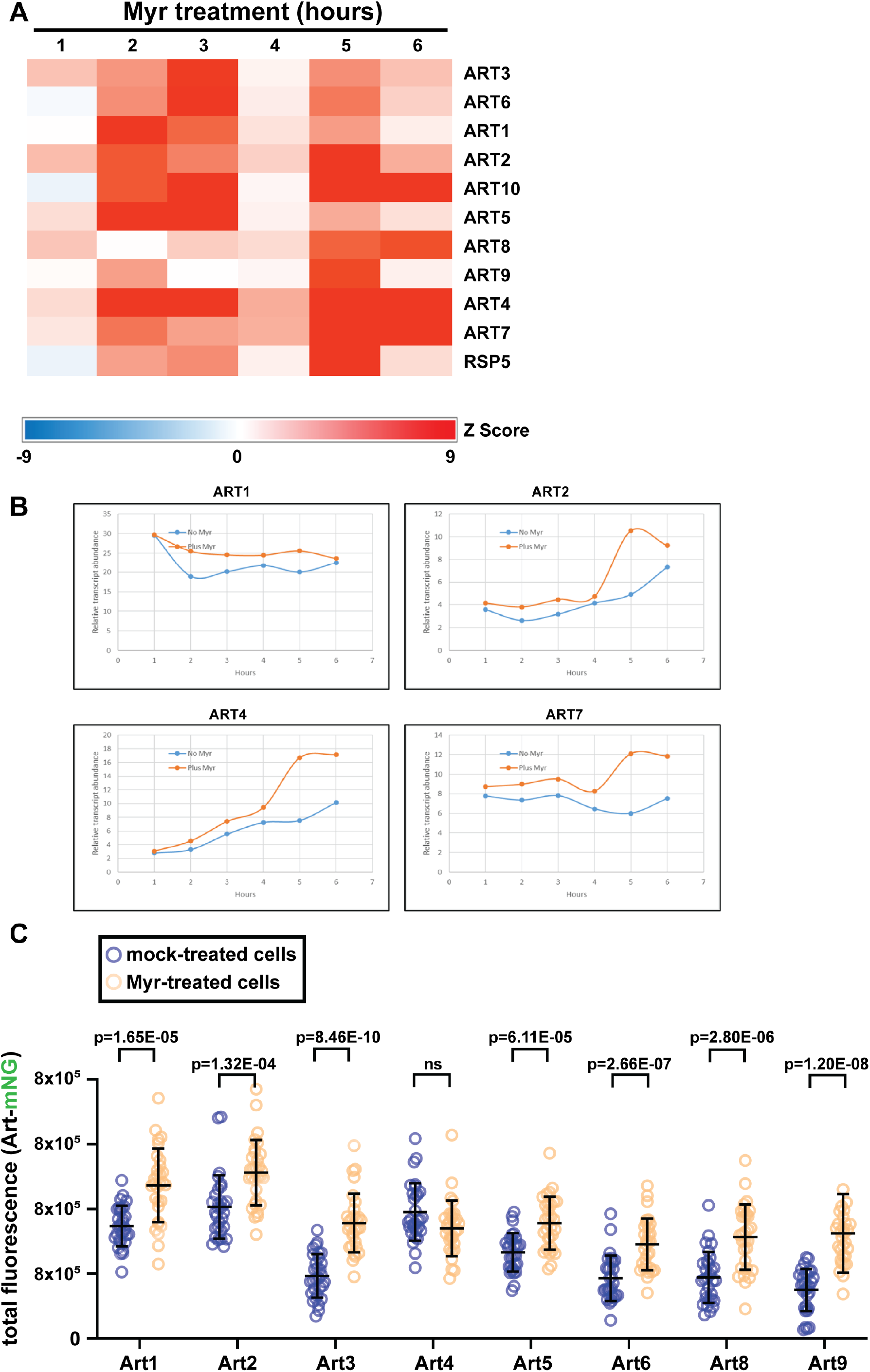
Myr increases the transcript and protein levels of most arrestin proteins. (A) Transcript levels of arrestins in Myr-treated cells as determined by RNAseq. (B) Transcript levels of ART1, ART2, ART4, and ART7 in cells treated with or without Myr as determined by RNAseq. (C) Myr increases the fluorescence of mNG-tagged arrestin proteins, except Art4.

**Figure S5.**
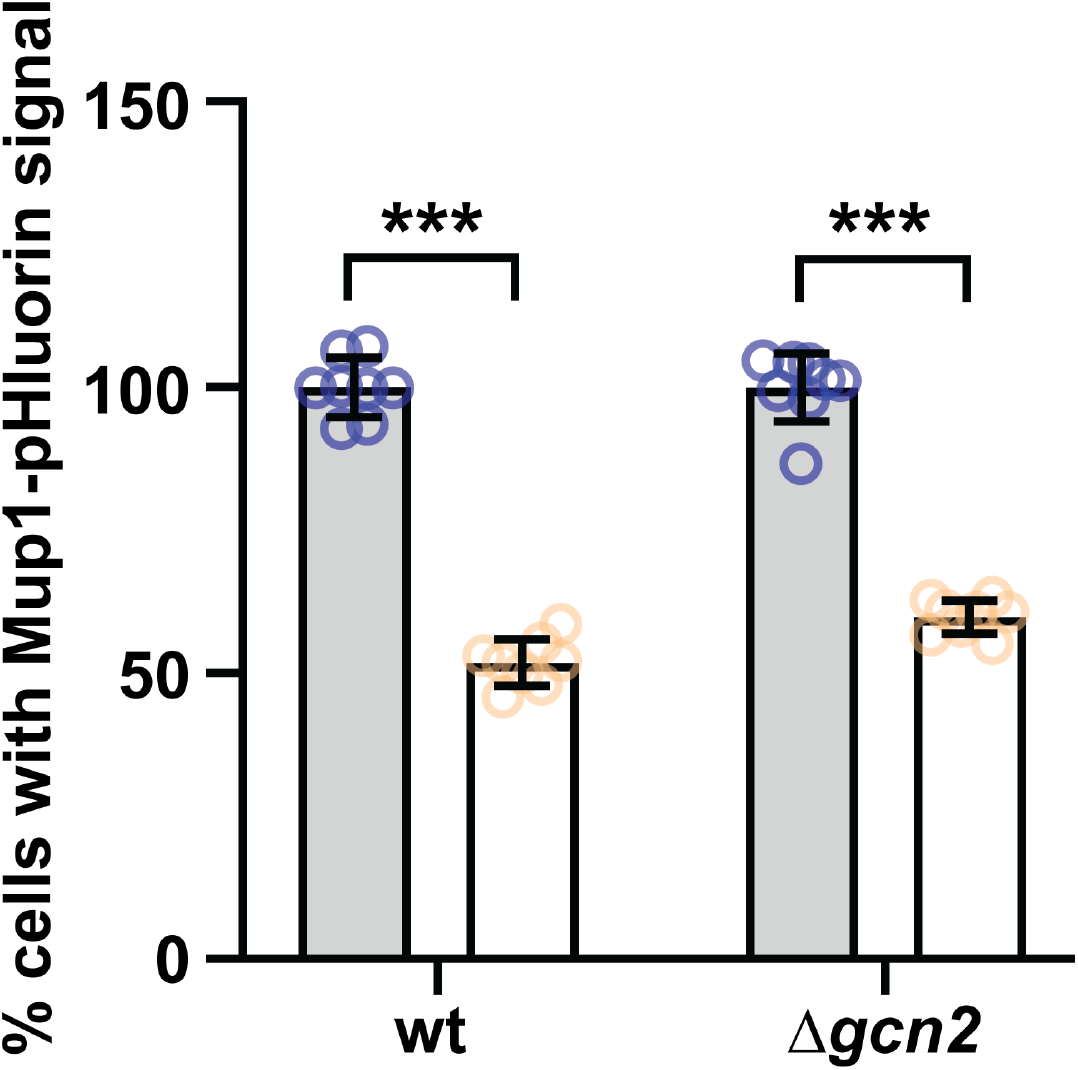
Myr-stimulated trafficking of Mup1 is Gcn2-independent. Flow cytometry analysis of wildtype or *Δgcn2* cells expressing Mup1-pHluorin in media with or without myriocin.

**Figure S6.**
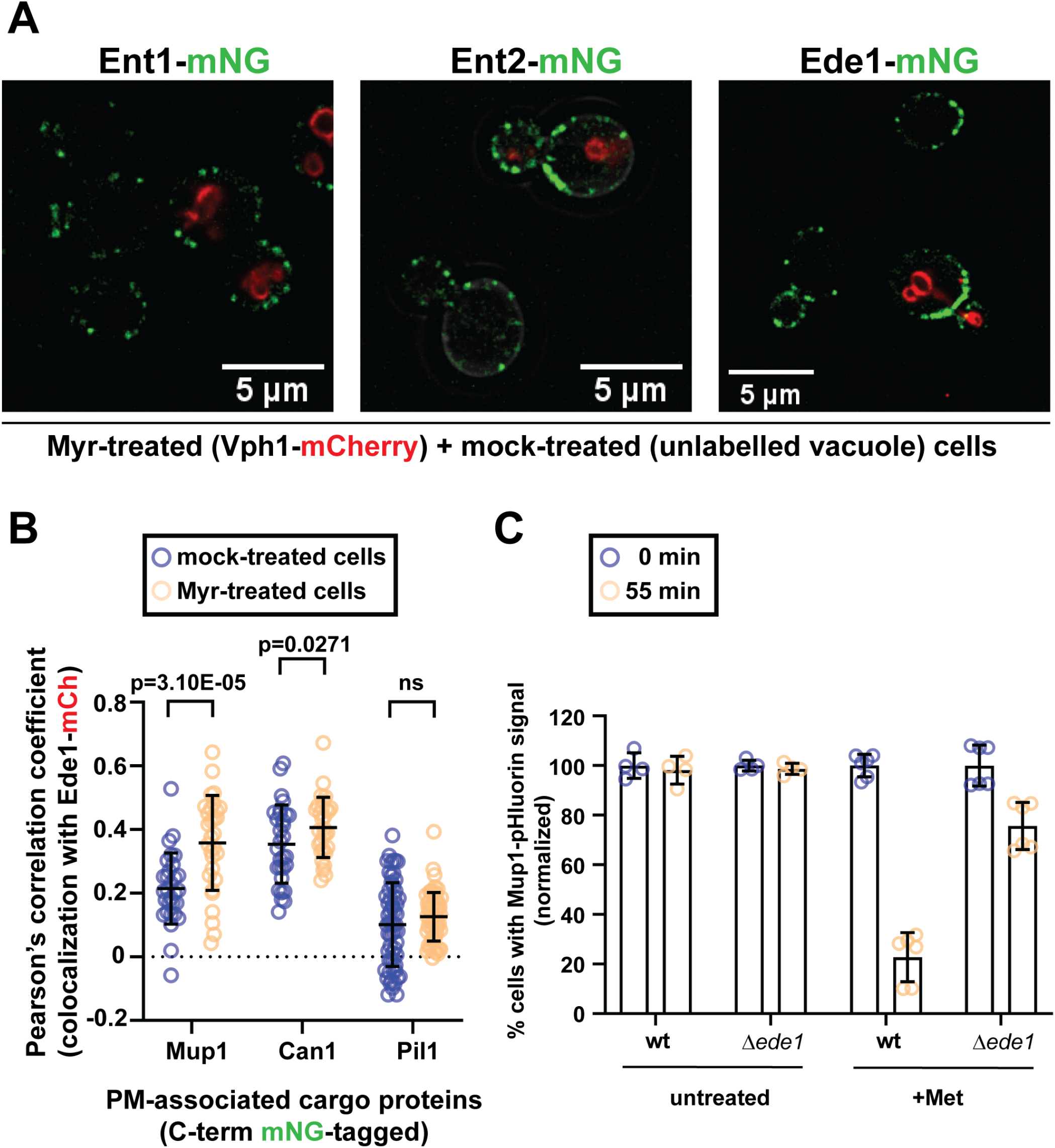
Analysis of endocytic adaptor localization in response to sphingolipid depletion. (A) Fluorescence microscopy of cells showing the increase of Ent1-, Ent2-, and Ede1-mNG fluorescence at the periphery of the plasma membrane after 5 hours of Myr treatment. (B) Colocalization of Ede1-mCh with select mNG-tagged PM integral proteins after 5 hours with or without Myr treatment. (C) Flow cytometry analysis of cells expressing Mup1-pHluorin in media with or without methionine.

## Acknowledgements

We are grateful to T Graham, A Ebert, and S Qualls-Histed for advice and helpful discussions. RCD was supported by NIH grant R56 AG024377 (to RCD). NLH and JAM were supported by NIH grant R35 GM144112 (to JAM).

## Author contributions

Conceptualization, N.L.H., R.C.D. and J.A.M.; Methodology, N.L.H., R.C.D. and J.A.M.; Investigation, N.L.H., B.M., R.C.D. and J.A.M.; Resources, R.C.D. and J.A.M.; Writing – Original Draft, N.L.H. and J.A.M.; Writing – Review & Editing, N.L.H., B.M., R.C.D. and J.A.M.

